# Spans attributed to short-term memory are explained by sensitivity to long-term statistics in both musicians and individuals with dyslexia

**DOI:** 10.1101/795385

**Authors:** Eva Kimel, Atalia Hai Weiss, Hilla Jakoby, Luba Daikhin, Merav Ahissar

## Abstract

Reduced short-term memory (STM) of individuals with dyslexia (IDDs) and enhanced STM of musicians are well documented, yet their causes are disputed. We hypothesized that their STMs reflect their sensitivities to accumulative long-term stimuli statistics. Indeed, when performing an STM task, IDDs had reduced benefit from syllable frequency, whereas musicians manifested an opposite effect, compared to controls. Interestingly, benefit from sequence-repetition did not significantly differ between groups, suggesting that it relies on different mechanisms. To test the generality of this separation across populations, we recruited a group of good-readers, whose native language contains a smaller fraction of the high-frequency syllables. Their span for these “high-frequency” syllables was small, yet their benefit from sequence-repetition was adequate. These experiments indicate that sensitivity to long-term stimuli distribution, and not to sequential repetition, is reduced in IDDs and enhanced in musicians, and this accounts for differences in their STM performance.

## The effective capacity of STM is sensitive to item frequency

The capacity of our short-term memory (STM) is limited, ranging from the classic estimation of 7 items (“magical number 7±2”, Miller, 1956) to the more recent suggestion of 4 items (“magical 4±1”, Cowan, 2010). The most common procedure for measuring STM is the Span Task, in which a participant is asked to reproduce a sequence of items in the order of their presentation. Many studies have found that the item characteristics greatly affect performance in these tasks. For example, spans of frequent words are larger than those of infrequent ones (Hulme et al., 1997; Roodenrys, Hulme, Lethbridge, Hinton, & Nimmo, 2002), and span for words is greater than for non-words (Hulme, Maughan, & Brown, 1991). The benefit of familiarity applies also at the syllabic level. Syllable spans are typically assessed with a slightly different task: non-word repetition (e.g. Gathercole & Adams, 1993 in 2-3 year old children; Susan E Gathercole & Baddeley, 1989; Susan E Gathercole, Frankish, Pickering, & Peaker, 1999). For example, Nimmo and Roodenrys (Nimmo & Roodenrys, 2002) showed that both non-word repetition accuracy and recall of syllable sequences are higher for syllables that occur more frequently in polysyllabic English words. Similarly, Tremblay and colleagues found that non-word repetition accuracy is greater when the frequency of their first syllable is high (Tremblay, Deschamps, Baroni, & Hasson, 2016).

## IDDs are considered to have reduced STM capacity and have reduced utilization of implicit prior knowledge

Developmental dyslexia is defined as a specific difficulty in “accurate or fluent word recognition, poor decoding, and poor spelling abilities” (American Psychiatric Association, 2013), in spite of adequate hearing levels, normal intelligence and adequate educational opportunities. Beyond reading difficulties, of individuals with dyslexia (IDDs) have substantial difficulties in phonological awareness and in span tasks (Jeffries & Everatt, 2004; Roodenrys & Stokes, 2001; Snowling, Goulandris, Bowlby, & Howell, 1986; Snowling, 1981, though see Wimmer, 1993).

Traditional theories of dyslexia attributed these difficulties to a core deficit in phonological representations (e.g. Snowling, 2000), or reduced efficiency in accessing these representations (Ramus, 2014; Ramus & Szenkovits, 2008). However, more recent theories, such as the “anchoring deficit hypothesis” (Ahissar, 2007; Ahissar, Lubin, Putter-Katz, & Banai, 2006) have proposed a learning-related core deficit: reduced sensitivity to stimulus statistics, in spite of adequate sensory processes (Jaffe-Dax, Raviv, Jacoby, Loewenstein, & Ahissar, 2015). Specifically, IDDs’ memory traces decay faster (Jaffe-Dax, Frenkel, & Ahissar, 2017; Jaffe-Dax, Kimel, & Ahissar, 2018; Lieder et al., 2019), yielding a functional “leakage” of memory traces, a consequent reduced learning rate, and hence poorer long-term representations (Banai & Ahissar, 2017). This hypothesis has some overlap with the phonological account – since phonological representations under this account are also expected to be somewhat impoverished. But, the predictions of this account are not specific to phonology, and in some cases it bears opposite predictions; learning-related deficit theories, as opposed to perceptually-based deficits, predict that relative difficulties are expected to increase with stimulus-specific exposure. Deficits associated with linguistic regularities, such as morphological deficits, have been reported in several studies (Rispens, Roeleven, & Koster, 2004; Schiff & Ravid, 2007). For example, IDDs have been shown to have a reduced morphological benefit in acquiring new words (Kimel & Ahissar, 2019). This learning deficit hypothesis leads us to expect an increase in relative difficulties with increased exposure, due to the reduced rate of learning of regularities and repetitions in the input. Given the evidence of substantial influence of item frequency on the performance in span tasks, we now ask whether IDDs’ reduced sensitivity to language statistics can explain one of IDDs’ most reliably reported characteristics – their reduced STM.

## Musicians are considered to have increased STM capacity and have enhanced utilization of implicit prior knowledge

There is ample evidence that musical proficiency is correlated with performance in STM tasks in the domain of language (Chen, Penhune, & Zatorre, 2008b, 2008a; Janata & Grafton, 2003; Janata, Tillmann, & Bharucha, 2002). For example, children and adults who have received musical training have been shown to outperform non-musicians on digit and non-word span tasks (Franklin et al., 2008; Fujioka, Ross, Kakigi, Pantev, & Trainor, 2006; Lee, Lu, & Ko, 2007; Parbery-Clark, Skoe, Lam, & Kraus, 2009, Chandrasekaran & Kraus, 2010).

In separate, seemingly unrelated studies, it has been found that musicians’ show enhanced performance in statistical learning tasks (Shook, Marian, Bartolotti, & Schroeder, 2013). We hypothesize that musicians’ enhanced STM is actually a manifestation of an improved utilization of long-term stimulus statistics, which beneficially impacts their ability to benefit from exposure.

## Sensitivity to stimulus statistics is expected to affect learning rate

We thus hypothesize that both IDDs’ poor STM and musicians’ elevated STM are a consequence of their relative sensitivities to long-term item statistics, specifically stimulus frequency. Figure 1 is a schematic illustration of this hypothesis and its predictions. The main underlying principle is that cumulative benefits from encounters with stimuli, i.e. learning rate, are affected by sensitivity to stimuli statistics (e.g. in the case of IDDs, sensitivity is reduced due to faster memory decay, and hence learning rate is decreased). Importantly, though learning rate decreases with exposure (learning curve is convex), it does not saturate, and improvement continues for years. This continuous learning has been shown across domains (the exponential/power law of practice – Crossman, 1959; Heathcote, Brown, & Mewhort, 2000), and, among others, applies to reading. Thus, although the reading rate of IDDs continuously improves with practice, they (typically) do not catch up with their peers. This basic conceptualization yields a simple, yet not intuitive prediction, as illustrated in Figure 1. Although all groups improve with practice, the group differences in performance will increase with greater practice and exposure (right hand side of Figure 1). Therefore, compared to controls with a similar exposure, IDDs’ STM will be particularly poor for frequent items, whereas musicians’ STM will be particularly high.

**Figure 1.**
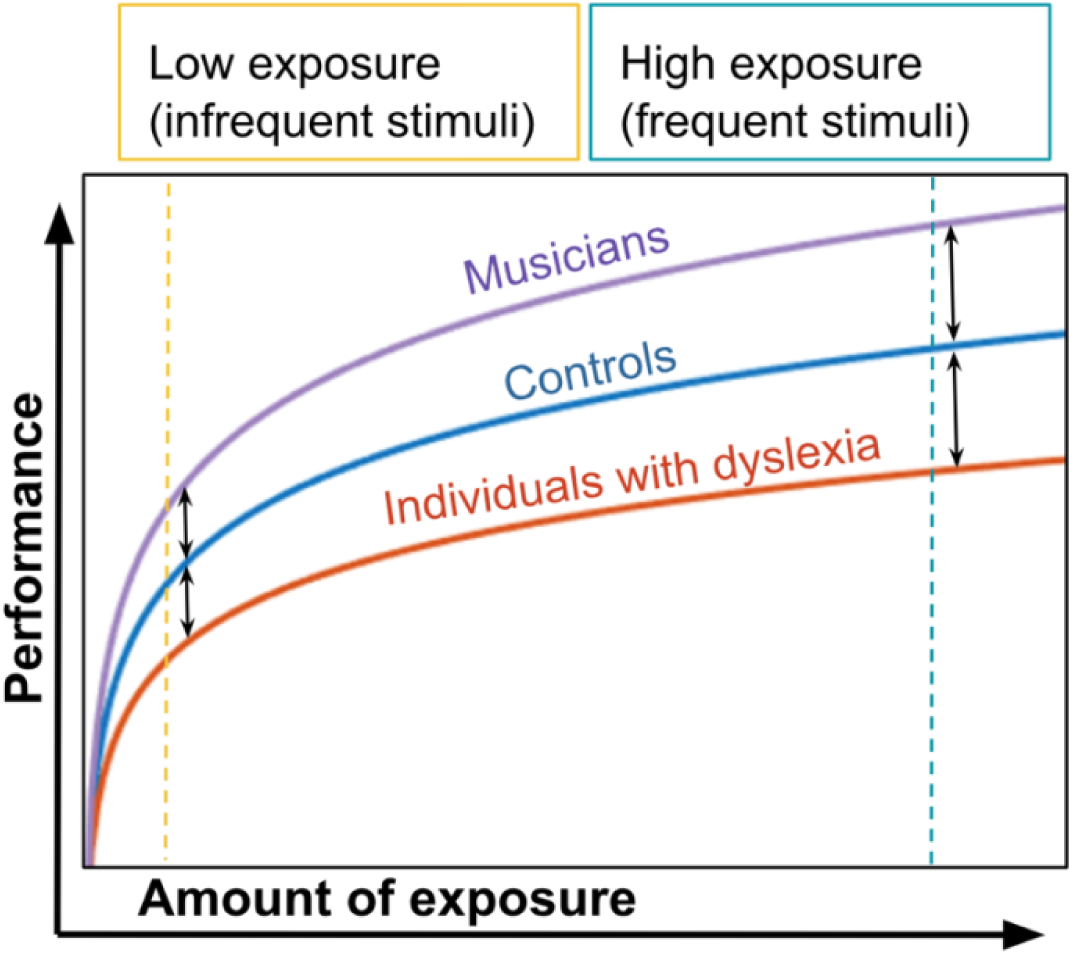
A schematic illustration of expected performance in span tasks as a function of exposure to the tested stimuli in 3 populations with different sensitivities to stimuli statistics. All participants start from scratch (zero performance) and improve, with no saturation (e.g. log function). Therefore, group difference is expected to increase with exposure, and be larger for high-frequency items (blue dashed line on the right) compared with low-frequency items (orange dashed line on the left). To test the impact of item frequency, we measured spans of two syllable types, frequent and infrequent, respectively.

As our stimuli we chose frequent and infrequent syllables. We chose to focus on syllables as their familiarity is not strongly dependent on reading experience; they usually do not carry semantic content in Hebrew; and there is evidence of their presence as separate mental entities quite early in development (e.g. Perfetti, Beck, Bell, & Hughes, 1987). Thus, illiterate adults and nursery-school children show similar success rates in manipulating syllables, but only literate individuals successfully operate with phonemes (Liberman, Shankweiler, Fischer, & Carter, 1974; Morais, Cary, Alegria, & Bertelson, 1979). As frequent syllables, we chose Consonant-Vowel (CV) syllables, and as infrequent we chose Vowel-Consonant (VC) syllables. CV syllables are part of the syllabic vocabulary in almost all known languages (Sommer, 1970). They are acquired early and predominate both in babbling and in early meaningful speech (Stoel-Gammon, 1989). In Hebrew they constitute the majority of syllables (Ben-Dror, Frost, & Bentin, 1995). Contrary to CV syllables, VC syllables are rare in languages in general (Clements & Keyser, 1983). In Hebrew VC syllables are an order of magnitude less frequent than CV syllables (Ben-Dror et al., 1995; discussed in Share & Blum, 2005).

## Experiment 1

We measured spans of frequent (CV) and infrequent (VC) syllables in 3 populations: IDDs, controls and musicians (selection criterions are detailed in the method section). Based on previous studies, mean spans were expected to be higher for frequent than infrequent syllables; overall performance of IDDs was expected to be lower than that of the control group; musicians’ performance was expected to be higher than controls’. Crucially, we predict a previously unattested interaction: syllabic frequency would benefit musicians more than controls, but aid IDDs to a lesser extent.

Span scores also benefit from sequence repetition (serial-order repetition). Importantly, some studies (Bogaerts, Szmalec, Hachmann, Page, & Duyck, 2015; Szmalec, Loncke, Page, & Duyck, 2011; though see Henderson & Warmington, 2017; Staels & Van den Broeck, 2014, 2015;Wang, Xuan, & Jarrold, 2016) have found that IDDs benefit less from serial-order repetition, and this might account for IDDs’ reduced spans (and hence also account for musicians’ increased spans). We therefore also assessed benefits from serial-order repetition in the same task. Assessing the specific advantage afforded by repetition of the same series, for each type of syllables (CV and VC) separately, provided a separate estimate for repeated series learning.

## Method

### General cognitive assessments

Non-verbal intelligence was assessed using the Block Design task (a subtest from the Hebrew version of the Wechsler Adult Intelligence Scale, WAIS-III; Wechsler, 1997). The Block Design task measures spatial reasoning abilities, and is often used to match groups on non-verbal reasoning.

Standard short-term memory skills were assessed with the sub-tests of Digit Span (Wechsler, 1997): Digit Forward and Digit Backward. Digit Forward requires immediate oral repetition of orally presented sequences of digits, and Digit Backward requires immediate oral repetition of orally presented sequences of digits in reversed order.

### Reading measures

We used three measures of reading proficiency: single word reading, pseudo-word reading, and paragraph reading. The lists of real words and pseudo-words were standard lists (Deutsch & Bentin, 1996), presented with diacritics, which make Hebrew orthography transparent. Reading in context was assessed by reading a four-paragraph academic-level text in Hebrew (standardized for students by our lab; Ben-Yehudah, Sackett, Malchi-Ginzberg, & Ahissar, 2001). Participants were instructed to read the text aloud, as quickly and accurately as possible, but slow enough to be able to answer a simple content question at the end, in order to encourage text comprehension. Both accuracy and rate (i.e. total number of words read in one minute) were scored for all reading tests.

### Stimuli – estimating syllabic frequency

The exact frequency of syllables in Hebrew has not been calculated, probably because syllables cannot be automatically parsed from written Hebrew without vowel diacritics. We therefore assessed syllable frequency by calculating the distribution of syllables in the largest publicly available corpus of spoken Hebrew that has a phonological transcript (~6,500 word tokens; The Corpus of Spoken Israeli Hebrew – CoSIH). In this corpus about 54% percent of the syllables are CV, 25% CVC, and only 5.5% VC. The mean (SD) % of occurrence in the corpus for the specific syllables that were used in the current study is 0.58 (0.69) for the CV syllables and 0.11 (0.11) for the VC syllables, t = 6.8, p < 10-8.

### Procedure

We used a Syllable Span protocol (Oganian & Ahissar, 2012; Weiss, Granot, & Ahissar, 2014) which is based on the Digit Span protocol (WAIS-III; Wechsler, 1997), and administered it four times: with frequent (CV) and infrequent (VC) syllables, with and without series repetition. VC syllables were constructed by switching the vowel and the consonant phoneme in each CV syllable, e.g. *ko* and *ok.* In the no-repetition condition, no syllable was repeated throughout the experiment. In the repeated condition we used two sequences that were gradually elongated and were presented in an interleaved manner. For example, the four initial sequences for the CV repeated condition were: **1.** /ve/ /tsu/ **2.** /ko/ /ʃa/ **3.** /ve/ /tsu/ /pi/ **4. /**ko/ /ʃa/ /ħu/; and for the non-repeated condition: **1. /**tsu/ /na/ **2.** /ʃe/ /ko/ **3. /**li/ /ʃu/ /ga/ **4.** /di/ /bu/ /mo/.

The VC spans were always administered before the CV spans, in order to avoid a possible benefit for the VC syllables due to training. The order of repetition vs. no-repetition was counter-balanced across participants. A recording of a native female Hebrew speaker was used; syllables were played at a fixed interval of 1 second (SOA). The score for each condition is the number of sequences which the participant reproduced correctly.

### Participants

Participants were recruited through ads posted at the Hebrew University campuses, two other colleges in Jerusalem, and the Jerusalem Academy of Music and Dance. Participants were paid for their participation. Candidate participants filled in a detailed questionnaire about their formal academic education, history of reading difficulties (including previous diagnoses), musical background and medical condition. Exclusion criteria were: hearing problems, psychiatric medications other than attention deficit medication, and formal musical background for control and IDDs (i.e. more than 2 years of either playing a tonal instrument or formal singing education). All participants received all their schooling in Israel. Individuals who passed this preliminary screening were invited to a basic assessment session of cognitive and reading skills. On the basis of their performance in this session, one participant with dyslexia and one control participant were excluded due to below average cognitive scores (i.e. Block Design score < 7; Wechsler, 1997). One IDD had accurate (100% correct) non-word reading, and was therefore excluded from the study. The dyslexic group included 8 participants who typically take medicine for ADHD (e.g., Concerta). Based on our previous studies (Ben-Yehudah et al., 2001) they were not excluded from participation, but they did not take medicine on test days.

Thirty-four control individuals, 35 IDDs and 37 musicians took part in Experiment 1. All members of the musician cohort were either students at the Jerusalem Academy of Music and Dance or professional musicians with an academic degree in music. Data of participants whose score was less than 3 in any of the span conditions were removed from the results, as we attribute their poor performance to lack of understanding of the task (n=1); the 3 oldest control participants and the 7 youngest musicians were excluded in order to match age with the other groups. Thus, we report results of 31 controls, 34 IDDs and 30 musicians matched for age and general reasoning, as shown in Table 1. As expected, there was a significant difference in reading and reading related measures. In addition, consistent with previous findings (e.g. Ackerman, Dykman, & Gardner, 1990; Torgesen, Wagner, Simmons, & Laughon, 1990), IDDs’ scores were significantly lower than those of controls and musicians in the Digit Span task.

**Table 1.**

|  | **Controls** | **IDDs** | **Musicians** |
| --- | --- | --- | --- |
| **Measure** | N = 31 (13 M) | N = 34 (11 M) | N = 30 (9 M) |
| **Years of musical practice** | Up to 2 | Up to 2 | 12.7 (3.7) |
| **Age (years)** | 25.16<sub>a</sub> (2.2) | 24.32<sub>a</sub> (3.1) | 24.27<sub>a</sub> (2.2) |
| **Block design (scaled)** | 12.71<sub>a</sub> (3.0) | 12.91<sub>a</sub> (2.7) | 13.73<sub>a</sub> (2.6) |
| **Digit Span (scaled)** | 11.32<sub>a</sub> (2.8) | 7.88<sub>b</sub> (1.9) | 12.13<sub>a</sub> (3.0) |
| **Single words reading accuracy (% correct)** | 96.91<sub>a</sub> (4.2) | 87.25<sub>b</sub> (8.0) | 99.17<sub>a</sub> (1.7) |
| **Single words reading rate (words/minute)** | 96.6<sub>a</sub> (29.5) | 69.27<sub>b</sub> (25.0) | 119.71<sub>c</sub> (22.5) |
| **Pseudo-words reading accuracy (% correct)** | 90.46<sub>a</sub> (11.2) | 60.78<sub>b</sub> (18.5) | 96.53<sub>a</sub> (3.6) |
| **Pseudo-words reading rate (words/minute)** | 57.82<sub>a</sub> (24.2) | 33.67<sub>b</sub> (11.9) | 76.26<sub>c</sub> (13.9) |
| **Paragraph reading accuracy (% correct)** | 98.65<sub>a</sub> (1.4) | 95.02<sub>b</sub> (4.3) | 98.85<sub>a</sub> (1.1) |
| **Paragraph reading rate (words/minute)** | 138.83<sub>a</sub> (21.0) | 98.69<sub>b</sub> (22.6) | 134.09<sub>a</sub> (15.9) |

Means (SD) are presented. M – Male participants.
Means with different superscript (a,b,c) significantly differ at the $p < 0.05$ level after Bonferroni correction.

## Results

The number of correctly reproduced sequences in each condition (score; Table 2) was analyzed using a mixed-design analysis of variance (ANOVA) with *Syllable Type* (frequent vs. infrequent) and *Repetition* (repeated vs. non-repeated) as within-subject factors, and *Group* (controls, IDDs, and musicians) as a between-subject factor.

**Table 2.**
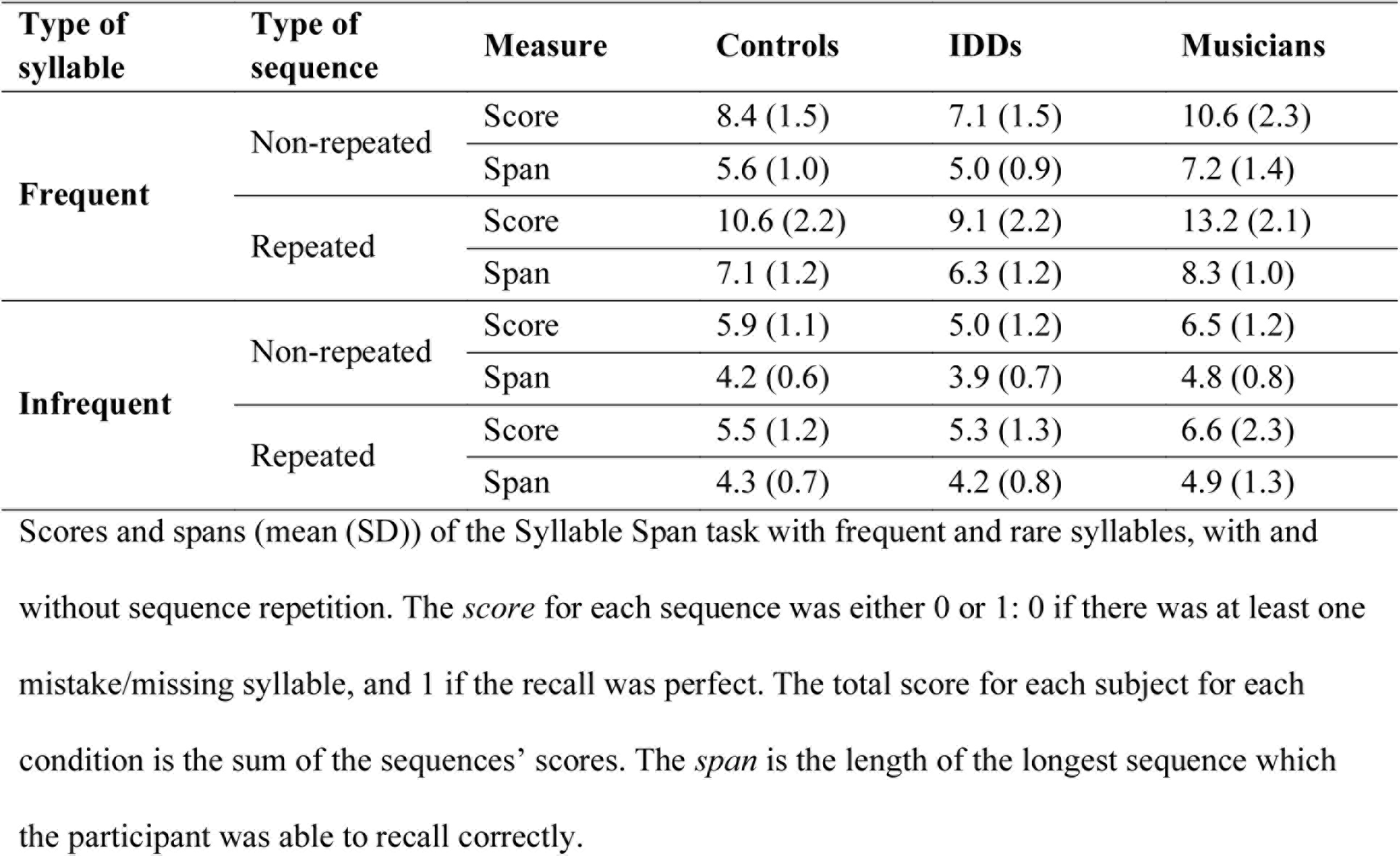

| Type of syllable | Type of sequence | Measure | Controls | IDDs | Musicians |
| --- | --- | --- | --- | --- | --- |
| Frequent | Non-repeated | Score | 8.4 (1.5) | 7.1 (1.5) | 10.6 (2.3) |
|  |  | Span | 5.6 (1.0) | 5.0 (0.9) | 7.2 (1.4) |
|  | Repeated | Score | 10.6 (2.2) | 9.1 (2.2) | 13.2 (2.1) |
|  |  | Span | 7.1 (1.2) | 6.3 (1.2) | 8.3 (1.0) |
| Infrequent | Non-repeated | Score | 5.9 (1.1) | 5.0 (1.2) | 6.5 (1.2) |
|  |  | Span | 4.2 (0.6) | 3.9 (0.7) | 4.8 (0.8) |
|  | Repeated | Score | 5.5 (1.2) | 5.3 (1.3) | 6.6 (2.3) |
|  |  | Span | 4.3 (0.7) | 4.2 (0.8) | 4.9 (1.3) |

Scores and spans (mean (SD)) of the Syllable Span task with frequent and rare syllables, with and without sequence repetition. The *score* for each sequence was either 0 or 1: 0 if there was at least one mistake/missing syllable, and 1 if the recall was perfect. The total score for each subject for each condition is the sum of the sequences' scores. The *span* is the length of the longest sequence which the participant was able to recall correctly.

The main effects were as expected: musicians achieved higher scores than controls, and controls achieved higher scores than IDDs (main effect of *Group*, F(2,92) = 33.18, p < 10-10, η2 =.419; a post hoc analysis with the Bonferroni correction: controls vs. IDDs: *p < .013*, controls vs. musicians: p *< 10-5*; Figure 2; Table 2). Scores for frequent syllables were higher than for infrequent ones (main effect of *Syllable Type*, *F*(1,92) = 762.3, *p <* 10-45, *η2 =* .892; Figure 2) in each of the groups (controls: *d =* 3.82, *SE =* .26, *p <* 10-25; IDDs: *d =* 3.0, *SE =* .25, *p <* 10-20, musicians: *d =* 5.37, *SE =* .26, *p <* 10-25).

**Figure 2.**
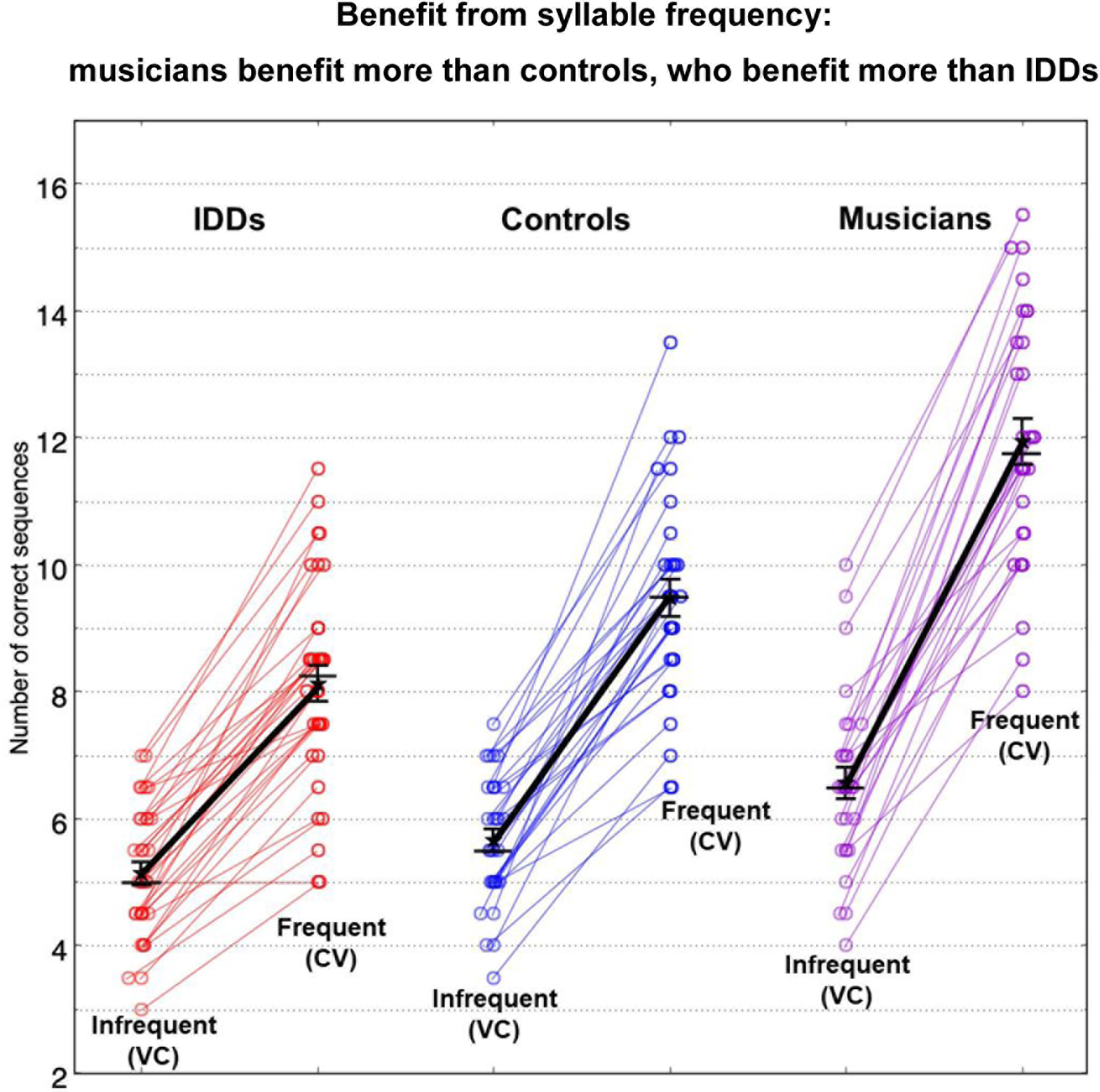
Scores of infrequent (VC) versus frequent (CV) syllable spans in the three test groups: IDDs (red), controls (blue), and musicians (violet). Groups are plotted (left to right) according to their increasing benefits from syllabic frequency. Spans are larger for frequent than for infrequent syllables (long-term frequency effect). Circles denote individual scores. Star symbols denote means; error bars denote 1 SEM; horizontal bars denote medians.

In line with our hypothesis, musicians benefited from syllable frequency more than controls, and controls benefited more than IDDs (*Syllable Type X Group* interaction, *F(2,92) = 22.2, p < 10-7, η2 =.325*; a one-tailed post hoc analysis on the difference between scores for frequent and for infrequent syllables with Bonferroni correction: controls vs. IDDs: *p < .034*, controls vs. musicians: p *< 10-4*; Figure 2). Additionally, group difference was larger for frequent (CV) than for infrequent (VC) syllables (CV syllables - Controls vs. IDDs: *d*=1.35, *SE*=.43, *p*<.007, Musicians vs. Controls: *d*=2.45, *SE*=.45, *p*<10-6, Musicians vs. IDDs: *d*=3.80, *SE*=.44, *p*<10-12; VC syllables - Controls vs. IDDs: *d*=.53, *SE*=.29, *p*=.203, Musicians vs. Controls: *d*=.91, *SE*=.29, *p*<.009, Musicians vs. IDDs: *d*=1.43, *SE*=.29, *p*<10-5; Bonferroni corrected for multiple comparisons).

Repeated sequences yielded higher scores than non-repeated sequences (main effect of *Repetition*, *F*(1,92) = 67.6, *p <* 10-11, *η2 =*.424; Figure 3; Table 2) in each of the groups (controls: *d =* .89, *SE =* .24, *p <* 10-3; IDDs: *d =* 1.15, *SE =* .23, *p <* 10-5, musicians: *d =* 1.37, *SE =* .25, *p <* 10-6). But in contrast to the group difference in benefits from syllable frequency, the three groups did not significantly differ in their benefits from sequence repetition (*Repetition X Group* interaction: *F*(2,92) = .98, *p =* .380, *η2 =*.021; Figure 3).

**Figure 3.**
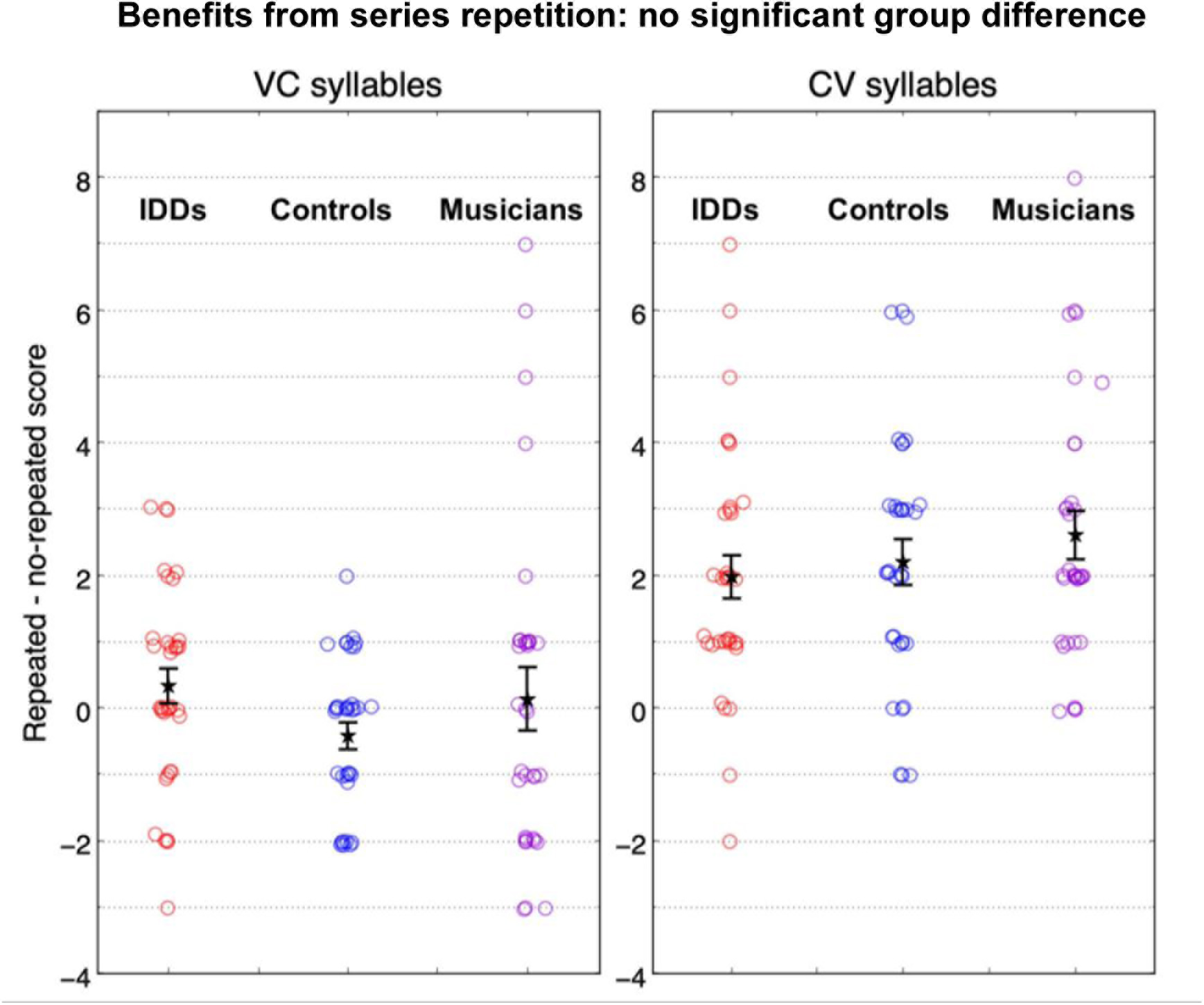
Difference between scores for spans with repeated series and spans without repeated series in sequences of infrequent (VC, left) and frequent (CV, right) syllables, in the three groups: IDDs (red), controls (blue), and musicians (violet). Benefit from series repetition does not significantly differ between the three populations, and is larger for CV syllables (right) than for VC syllables (left). Circles denote individual scores. Star symbols denote means; error bars denote 1 SEM; horizontal bars denote medians.

The benefit from repetition was larger for the frequent CV than for infrequent VC syllables (Syllable *Type X Repetition* interaction: *F*(1,92) = 67.1, *p <* 10-11, *η2 =*.422). In fact, repetition effect was significant for the frequent (*d =*2.26, *SE =*.20, *p <* 10-18), but not the infrequent syllables (*d =*.01, *SE =*.19, *p =* .947). Overall, the pattern of the benefit from repetition as a function of syllable frequency did not significantly differ between the groups (*Syllable Type X Repetition X Group* interaction: F(2,92) = 1.25, p < .291, η2 =.027; controls: benefit from repetition for frequent syllables *d =*2.19, *SE =*.35, *p <* 10-7, and for infrequent *d =*-0.42, *SE =*.33, *p =* .207; IDDs: benefit from repetition for frequent syllables *d =*1.97, *SE =*.33, *p <* 10-7, and for infrequent *d =*.32, *SE =*.32, *p =* .307; musicians: benefit from repetition for frequent syllables *d =*2.60, *SE =*.36, *p <* 10-10, and for infrequent *d =*.13, *SE =*.34, *p =* .692).

## Discussion

Overall the three groups showed the expected larger spans for lists composed of frequent (CV) vs. those composed of infrequent (VC) syllables, but IDDs’ gain from syllable frequency was smaller than controls’, and musicians’ gain was greater than controls. Our results, taken together with the typical use of frequent stimuli in STM assessments, suggest that the reports on musicians’ enhanced STM and IDDs’ reduced STM stem to a large extent from musicians’ enhanced and IDDs’ reduced use of long-term statistics. By contrast, the benefits of IDDs and musicians from sequence repetition did not significantly differ from controls’, suggesting that the benefits from series repetition do not differ between groups, and do not explain the difference in spans.

The group differences in item frequency benefit, together with their similar benefit from sequence repetition, suggest a dissociation between benefit from long-term item frequency and learning of a repeated sequence. In all three groups the benefit from sequence repetition was larger for CV than for VC sequences. This difference seems puzzling - if benefit from serial-order repetition does not depend on item familiarity, as suggested from the no group-difference in serial-order repetition, what other difference underlies the larger advantage of CVs compared with VCs in sequence repetition? A likely candidate is the “chunkability” of these two types of syllables, associated with their different manner of articulation, rather than their different degrees of familiarity. Thus it is easier to join the CV syllables to “chunks”, increasing the effective STM (Miller, 1956). More formally, CVs are categorized as “light syllables” whereas VC are categorized as “heavy syllables” (Duanmu, 2010; Lunden, 2011), based on their different type of ending (rime). Heavy syllables attract stress (the Weight-Stress Principle, Duanmu, 2010; Velupillai, 2012) and a word (or a chunk of syllables) can only have one primary stress. Therefore, a sequence of VCs tends to be perceived and articulated with a stress on each syllable, supporting segmentation rather than chunking, since stressed syllables are treated as word onsets (Cutler & Norris, 1988). By contrast, a sequence of CVs can be perceived and articulated as a long (chunked) multi-syllabic word.

Based on the results of Experiment 1, we propose that there is a group difference in sensitivity to the frequency of syllables, but there is no significant group difference in the sensitivity to series repetition, since effective learning of repeated series relies mainly on syllable “chunkability”, and the ability to chunk does not differ between the groups. To test this hypothesis, we conducted Experiment 2. The aim of this experiment was to test the dissociation between benefits from syllable frequency and syllable “chunkability” in a group that has adequate sensitivity to frequency, but reduced familiarity with the CV syllables. For this group we predicted reduced spans, but similar benefits from series repetition.

## Experiment 2

CV syllables constitute the majority of syllables in Hebrew (Ben-Dror, Frost, & Bentin, 1995), but their prevalence in English is substantially lower (28-38%, Dauer, 1983). Based on this substantial difference, we now hypothesized that native English speakers (whose familiarity with Hebrew is only basic) would show reduced CV spans compared with native Hebrew speakers (as illustrated in Figure 1 – left side vs. right side). We further hypothesized that, in spite of their reduced spans, their benefits from serial-order repetition of these CV syllables would be similar to those of native Hebrew speakers, since their “chunkability”, underlying the main benefits from repetition, is not based on familiarity but on their manner of articulation. We reasoned that such a dissociation would directly indicate the lack of dependence of benefits from series repetition on familiarity with the composing items, contrasting it with the dependence of STM on familiarity with the test items.

## Method

### Participants

Native English speakers without reading difficulties were recruited through ads put at the school of international students of the Hebrew University. All recruited participants took Hebrew as a second language class, and their familiarity with Hebrew was only basic. We applied similar inclusion criteria as for the Hebrew-speaking controls (e.g. no learning difficulties and musical education of up to 2 years). Additionally, the data of the youngest English speaking participant were removed in order to match age across the groups. Thus, the control group of Experiment 1 and the native English speakers groups were matched for age, general reasoning skills (based on the same criteria) and scaled Digit Span scores in their native language (Table 3).

**Table 3.**

| Measure | Hebrew-speaking Controls | English-speaking Controls | t-value |
| --- | --- | --- | --- |
|  | N = 31 (13 M) | N = 29 (15 M) |  |
| Age (years) | 25.2 (2.2) | 23.6 (2.2) | t = 1.55 ^ |
| Block design (scaled) | 12.7 (3.0) | 11.6 (2.6) | t = 1.64 ^ |
| Native language Digit Span (scaled) | 11.3 (2.8) | 11.6 (3.0) | t = -0.43 ^ |

Means (SD) are presented. M – Male participants. ^ - n.s.

Data of participants whose score was < 3 in one of the span conditions were removed from the results due to our attribution of their poor performance to misunderstanding of the task; data of one English speaking participant was consequently excluded. Participants for whom the difference between the conditions with and without repetition was smaller than their group mean by more than 2.5 standard deviations were defined as outliers, and were removed from the analysis; one additional English speaking participant was consequently excluded (for this participant the difference was −4). Thus, we report the results of 29 English speaking controls.

### Procedure

We used the same procedure as in Experiment 1, and administered it twice using CV syllables with and without repetition (Oganian & Ahissar, 2012; Weiss et al., 2014). The order of repetition vs. no-repetition was counter-balanced across participants. In order to have English-like syllables, a recording of a native female English speaker with native English articulation was used.

## Results

The number of correctly reproduced sequences in each condition (score) was analyzed using a mixed-design analysis of variance (ANOVA) with *Syllable Frequency* (frequent vs. infrequent) and *Repetition* (repeated vs. non-repeated) as within-subject factors, and *Group* (controls, IDDs, musicians, and English speaking controls) as a between-subject factor. For Hebrew-speaking controls, IDDs and musicians, a subset of the data collected in Experiment 1 (CV syllables only) is reanalyzed in Experiment 2.

As expected, scores of English-speaking controls for the CV syllables were lower than those of Hebrew speaking controls, in spite of their matched Digit Span scores in their native language (Table 3). Their CV spans were even lower than Hebrew speaking IDDs’ (main effect of *Group*: *F*(3,120) = 55.73, *p <* 10-21, *η2 =*.582; Bonferroni Post Hoc tests: IDDs vs. English speaking controls: *d =*1.51, *SE =*.42, *p <* .003; Figure 4). As expected, scores for repeated sequences were higher than for non-repeated (main effect of *Repetition*: *F*(1,120) = 154.17, *p <* 10-22, *η2 =*.562; Figure 4). However, there was no difference in the effect of repetition between the groups (*Repetition X Group*: *F*(3,120) = 1.76, *p =* .159, *η2 =*.042; Bonferroni Post Hoc tests were all non-significant, for example, musicians vs. English speaking controls: *d =*1.08, *SE =*.48, *p =* .160; Figure 4).

**Figure 4.**
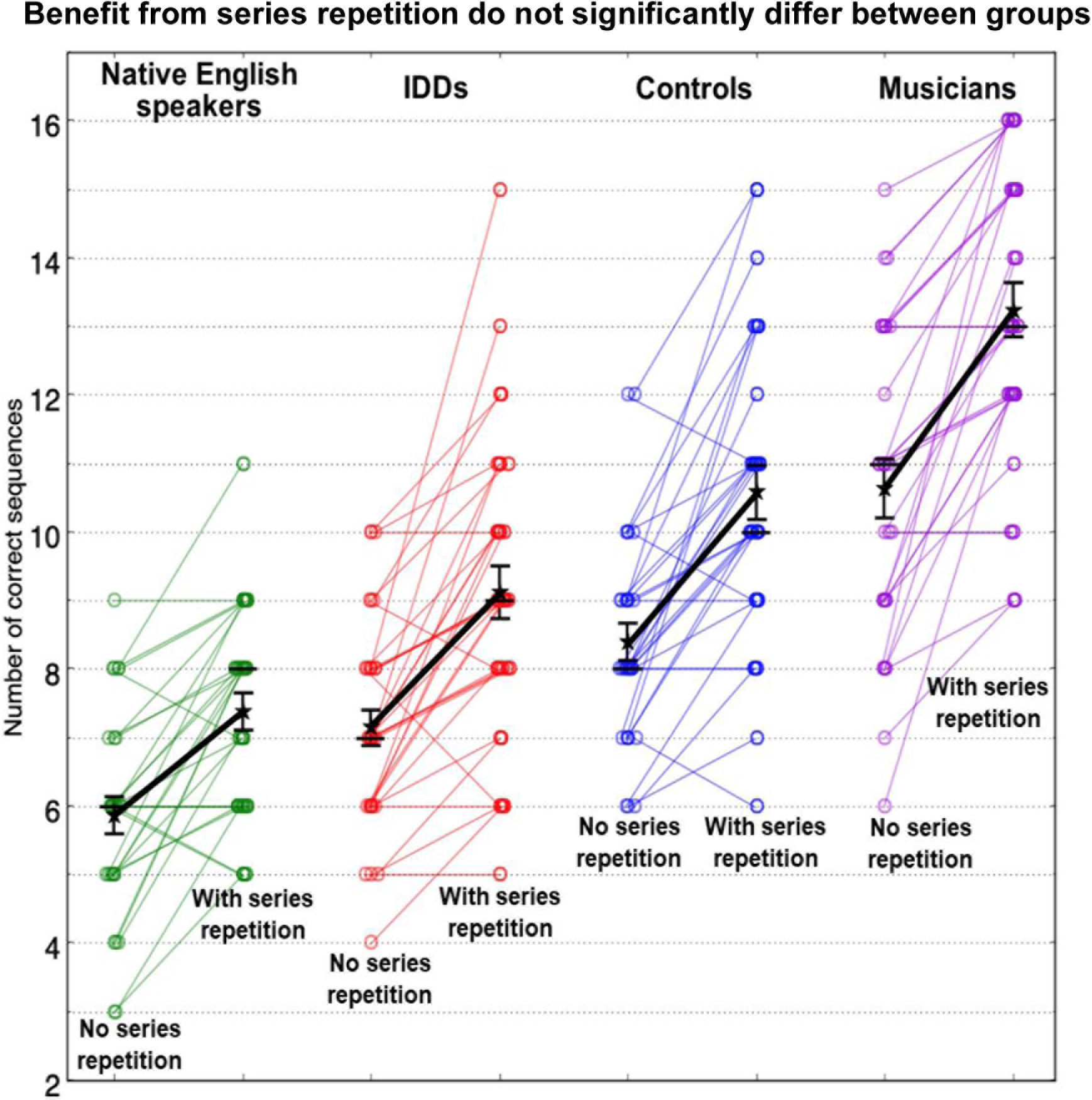
Spans of CV syllables, with and without series repetition: 4 populations with different degrees of exposure (Hebrew versus English speakers) and different sensitivity to item frequency (IDDs, controls. and musicians) differ in absolute scores but not in the benefits from series repetitions. CV span scores of native English speakers (green), native Hebrew-speaking IDDs (red), controls (blue), and musicians (violet). For each participant: left - without repetition, and right - with repetition. Spans of musicians are larger than controls’, which are larger than IDDs’. Spans of all Hebrew speakers are larger than of English speakers. However, benefits from series repetition do not significantly differ between the groups. Circles denote individual scores. Star symbols denote means; error bars denote 1 SEM; horizontal bars denote medians.

## Discussion

As predicted, native English speakers, who are less familiar with CVs, had smaller CV spans than native Hebrew speakers. Yet, they did not significantly differ from Hebrew speakers in their benefit from series repetition of these syllables. These results support the hypothesis that the benefits due to series repetition, do not substantially rely on item frequency, beyond basic familiarity - the frequency of CV syllables is lower for native English speakers, and still their benefit from sequence repetition is comparable to native Hebrew speakers’.

### General Discussion

In Experiment 1, we measured recall spans for infrequent (VC) syllables, and compared them to spans for frequent (CV) syllables, with and without sequence repetition in three populations: Hebrew-speaking IDDs, controls and musicians. IDDs benefited from syllable frequency less than controls, and musicians benefited more than controls. However, the benefit in span for serial-order repeated series did not significantly differ between the groups. To further separate the factors underlying the benefit of familiarity from the factors underlying the benefit of serial-order repeated series we conducted Experiment 2. Native English speakers, less familiar with CV syllables, were administered the CV span task, with and without sequence repetition. Their scores were very poor, as expected. Yet, they showed a similar benefit from sequence repetition as was seen in Hebrew speakers.

The group difference in the benefit from syllable frequency sheds light on the mechanisms underlying IDDs’ reduced, and musicians’ enhanced, spans. Span tasks are usually performed using frequent items, and thus, it is likely that reports on reduced performance of IDDs and enhanced performance of musicians in fact rely to a large extent on reduced/enhanced benefit from item frequency. These results challenge the interpretation of span tasks as a basic measure of STM, and its implication that IDDs’ reduced spans, and musicians’ enhanced spans, indicate some unique STM properties. Importantly, they agree with recent imaging results suggesting that STM does not have representations separately from long-term memory, but rather reflects access to long-term memory (Sreenivasan, Curtis, & D’Esposito, 2014).

By contrast, we found no significant difference in the benefit from series repetition between the groups. In the literature, the impact of sequence repetition on IDDs’ performance is disputed (see Majerus & Cowan, 2016 for a review). For example, a Hebb repetition learning paradigm (Hebb, 1961), in which a participant is exposed to a repeated sequence of syllables interleaved with non-repeated sequences, was previously administered to adult IDDs. While some studies reported that sequence learning poses a unique difficulty in dyslexia (Bogaerts et al., 2015; Szmalec et al., 2011), others failed to replicate this observation, in spite of using the same stimuli and analyses (Staels & Van den Broeck, 2014, 2015). Our results suggest no deficit in serial-order learning among IDDs.

It has been previously suggested that musicians’ enhanced STM might rely on enhanced general chunking skills, which are developed as part of the musical training (Talamini, Altoè, Carretti, & Grassi, 2017). However, the generalization of musical training to enhanced chunking ability of untrained items is conceptually questionable given the specificity of benefits from training (Jakoby, Raviv, Jaffe-Dax, Lieder, & Ahissar, 2019). Still, enhanced chunking skills may be associated with innate musical competence, or other characteristics of individuals who become musicians (Corrigall, Schellenberg, & Misura, 2013). Given the lack of group difference in benefits from series repetition, our results do not support enhanced chunking in musicians.

Taken together, our results propose mirror skills of musicians and IDDs – enhanced versus reduced accumulation of stimuli statistics. Musician’s enhanced reading skills have been characterized before (e.g. Weiss et al., 2014; early reading in children Anvari, Trainor, Woodside, & Levy, 2002), and as such seem to mirror those of IDDs. However, the current study is the first that proposes, and shows, that for both populations, their unique STM characteristics can be explained by the same underlying mechanism, which affects their long-term representations.

Overall, a significant group difference in the benefit from item frequency alongside no significant group difference in the benefit from series repetition, suggest that these are driven by separate mechanisms. However, the two mechanisms interact following a large number of repetitions of a sequence. With abundant exposure, the repeated sequence becomes a new independent item (a stable long-term memory trace), which in turn can be used as part of a new sequence with other items (Reder, Paynter, Diana, Ngiam, & Dickison, 2007), and enhance STM, which relies on extensive long-term knowledge (e.g. Nimmo & Roodenrys, 2002). We found a significant group difference in the benefit from item frequency, but no significant group difference in the benefit from sequence repetition. Perhaps, manipulation of sequences does not differ between the groups, but when they are integrated and stored as independent units, the robustness of their representation, or the access to these representations, is reduced among IDDs and is enhanced among musicians.

## Acknowledgements

This work was supported by the Canadian Institutes of Health Research, the International Development Research Center, the Israeli Science Foundation, and the Azrieli Foundation (grant No. 2425/15), and a personal grant from the Israel Science Foundation (grant No. 1650/17). Declarations of interest: none.

